# A history of repeated antibiotic usage leads to microbiota-dependent mucus defects

**DOI:** 10.1101/2024.03.07.583875

**Authors:** Kertu Liis Krigul, Rachel H. Feeney, Supapit Wongkuna, Oliver Aasmets, Sandra M. Holmberg, Reidar Andreson, Fabiola Puértolas-Balint, Kateryna Pantiukh, Linda Sootak, Tõnis Org, Tanel Tenson, Elin Org, Bjoern O. Schroeder

**Author notes:** Corresponding authors: Elin Org, Riia 23b, 51010 Tartu, Estonia, Bjoern O. Schroeder, Försörjningsvägen 2A, 90187 Umeå, Sweden. These authors contributed equally (Kertu Liis Krigul, Rachel H. Feeney). These authors contributed equally (Elin Org, Bjoern O. Schroeder).

## Abstract

Recent evidence indicates that repeated antibiotic usage lowers microbial diversity and lastingly changes the gut microbiota community. However, the physiological effects of repeated – but not recent – antibiotic usage on microbiota-mediated mucosal barrier function are largely unknown. By selecting human individuals from the deeply-phenotyped Estonian Microbiome Cohort (EstMB) we here utilised human-to-mouse faecal microbiota transplantation to explore long-term impacts of repeated antibiotic use on intestinal mucus function. While a healthy mucus layer protects the intestinal epithelium against infection and inflammation, using *ex-vivo* mucus function analyses of viable colonic tissue explants, we show that microbiota from humans with a history of repeated antibiotic use causes reduced mucus growth rate and increased mucus penetrability compared to healthy controls in the transplanted mice. Moreover, shotgun metagenomic sequencing identified a significantly altered microbiota composition in the antibiotic-shaped microbial community, with known mucus-utilising bacteria, including *Akkermansia muciniphila* and *Bacteroides fragilis*, dominating in the gut. The altered microbiota composition was further characterised by a distinct metabolite profile, which may be caused by differential mucus degradation capacity. Consequently, our findings suggest that long-term antibiotic use in humans results in an altered microbial community that has reduced capacity to maintain proper mucus function in the gut.

## INTRODUCTION

The colon is densely populated by commensal bacteria that play an important role in aiding digestion and preventing colonisation by pathogenic bacteria. However, if not controlled by the host, the intestinal bacteria can pose a threat. Therefore, a gel-like mucus layer, which lines the colonic epithelium, acts as barrier to both the native bacteria and any potential pathogens by physically separating the epithelium from the luminal content. Moreover, while initially considered to be only a lubricant for faecal material, the mucus layer has since been shown to play an important role in host immune defence and immune cell signalling regulation, as well as acting as a reservoir for signalling peptides (Aihara et al., 2017).

A healthy mucus layer can be described as having a gradient, consisting of a sterile and densely structured inner area, lying closest to the epithelium, and a more loosely structured outer area that is colonised by the native microbiota and periodically washed away along with faecal material. To replenish the layer, mucus is continuously secreted by goblet cells located in the epithelium thereby creating a luminal-directed flow, referred to as the mucus growth rate, which is estimated as ∼2µm/min in healthy mice and ∼4µm/min in healthy humans (Birchenough et al., 2016; Gustafsson et al., 2012). This mucus growth actively maintains a safe distance between the epithelium and the microbiota.

The presence of the gut microbiota is known to be crucial for the development of a functioning mucus layer, as illustrated in germ-free mice where the lack of microbiota results in a dysfunctional, penetrable inner mucus layer (Johansson et al., 2015). Moreover, the composition of the gut microbiota can affect mucus function and development (Jakobsson et al., 2015). A change in composition can result in certain bacterial species beginning to degrade the mucus layer at an excessive rate, contributing to altered mucus structure which allows bacteria to move closer to the epithelium, consequently triggering inflammation.

Recent studies have shown that changes in gut microbiota composition, for example through consumption of a low-fibre diet, caused defects in the colon mucus layer (Desai et al., 2016; Schroeder et al., 2018). Moreover, impaired mucus function has been observed in colitis, diabetes and obesity in mice (Johansson et al., 2014; Miranda et al., 2019; Schroeder et al., 2020; Shen et al., 2019) and has been connected to metabolic disease (Chassaing et al., 2017) and ulcerative colitis (Johansson et al., 2014; Van Der Post et al., 2019) in humans. Interestingly, the mucus defect may even precede the onset of ulcerative colitis and thereby contribute to its development (Van Der Post et al., 2019).

Besides diet, antibiotic intake has a major impact on gut microbiota composition (Anthony et al., 2022; Blaser, 2016). By assessing data collected from volunteers in the Estonian microbiome cohort (EstMB) we could recently show that a history of repeated antibiotic usage caused perturbations to the gut microbiota, including loss of diversity, that can persist over time (Aasmets et al., 2022). With the dramatic rise in global antibiotic use in recent years, an increase in various complex diseases, including asthma (Korpela et al., 2016), allergies (Hirsch et al., 2017), celiac disease (Mårild et al., 2013), diabetes (Livanos et al., 2016) and chronic inflammatory diseases of the gastrointestinal tract (Fenneman et al., 2022), has been observed. Interestingly, a maternal antibiotic-perturbed microbiota has been shown to exacerbate gut inflammation when transferred to mouse pups deficient in the anti-inflammatory cytokine interleukin 10 (IL10^-/-^) (Schulfer et al., 2017), and antibiotic usage is significantly associated with an increased risk of new-onset IBD (Duan et al., 2024), indicating a connection between antibiotic use, microbiome composition and gut health.

While our previous work revealed the significant impact of antibiotic usage on microbiome composition and links between microbiota changes and disease (Aasmets et al., 2022), little is known about the long-term impact of antibiotic treatment on mucosal barrier function. As antibiotics can modify microbiota composition and the microbiota can, in turn, affect mucus characteristics, we hypothesised that antibiotic usage may lead to changes in the colonic mucus layer. To address this, we utilised our deeply phenotyped EstMB and selected human microbiota samples from individuals with a history of repeated, but not recent, antibiotic usage and a matched control group. Using human-to-mouse faecal microbiota transplantation (FMT) alongside *ex-vivo* mucus function assessment from viable tissue, we then studied the impact of antibiotic-altered human microbiota on the mouse gut environment, with a particular focus on the colonic mucus layer.

## MATERIALS AND METHODS

### Human donor recruitment and sample collection

Human donors were recruited as part of the Estonian Microbiome (EstMB) project established in 2017, as described in detail previously (Aasmets et al., 2022). Briefly, 2509 volunteers from the Estonian Biobank, which contains more than 210,000 genotyped adults, donated stool, oral and plasma samples for the microbiome study. The cohort currently includes 1764 female and 745 male participants, aged 23-89 years. The participants collected a stool sample immediately after defecation and delivered it to the study centre, where it was stored at −80 °C until further processing.

All participants provided informed consent and signed a broad consent form, which allows access to the participant’s personal and medical health records data from national health registries and databases. Additionally, the patients reported their health-related behaviour by completing a lifestyle questionnaire, which included questions about their diet (i.e. food frequency questionnaire (FFQ)), physical activity, medical data, living environment, and stool characteristics (i.e. Bristol stool scale).

The study was approved by the Research Ethics Committee of the University of Tartu (approval No. 266/T10) and by the Estonian Committee on Bioethics and Human Research (Estonian Ministry of Social Affairs; approval No. 1.1-12/17). The rights of gene donors are regulated by the Human Genes Research Act (HGRA) § 9 – Voluntary nature of gene donation.

### Human donor inclusion criteria

The following inclusion criteria were used for this study: Healthy adults of both sexes with normal BMI (between 18.5 - 25), no medical history of complex diseases (listed in **Supplementary Table 1**) and had not taken any medications in the 3 months prior to stool sample collection (listed in **Supplementary Table 2**) nor antidepressants in the 5 years prior to stool sample collection. Additionally, participants who had seasonal allergies or celiac disease according to the questionnaire data were excluded.

The donor group of repeated antibiotics users (hABX) did not take any antibiotics in the 6 months prior to stool sample collection but had used antibiotics at least 5 times within the last 5 years. For the control group of human donors (hCTRL), individuals who had not used antibiotics in the last 10 years were included. Vegans and vegetarians were excluded from the set of cases and controls to avoid any diet-dependent bias on microbiota composition. When selecting matched controls, the following factors were considered: age, sex, BMI, Bristol stool scale (stool type) and diet. In summary, from 2509 individuals, four cases were selected which were matched with four controls (**Figure 1**).

**Figure 1.**
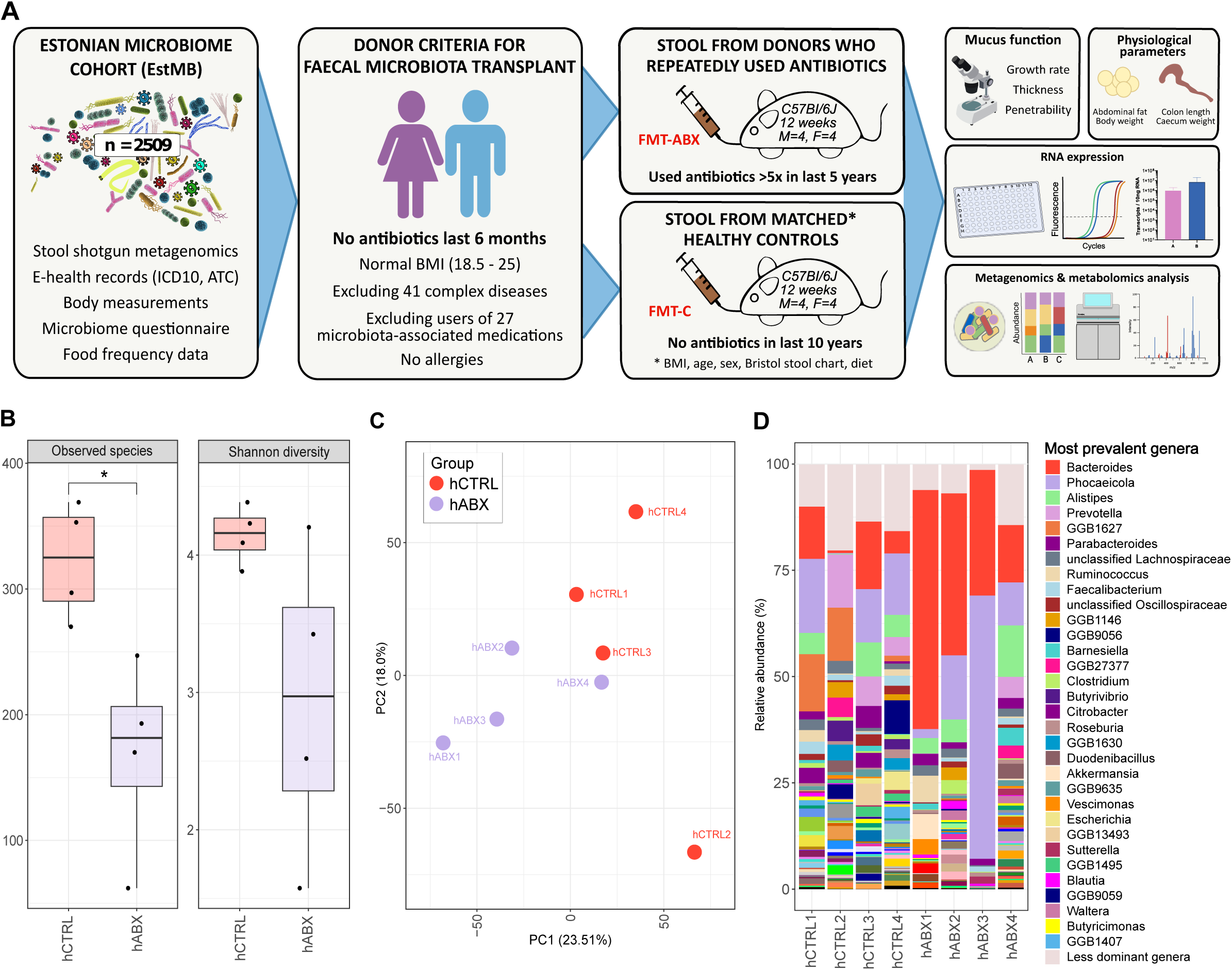
Overview of the donor selection, experimental plan and donor microbiomes. (A) Workflow of faecal microbiota donor selection from the EstMB participants, including mouse sample collection and analysis (created with biorender.com); (B) Alpha diversity analyses (observed number of species and Shannon diversity index) of individual human donor stools (hCTRL, hABX); (C) Beta diversity of the individual human donor stools and pooled donor stool; (D) Relative abundance of most common genera in hCTRL and hABX individual donors; hCTRL - human controls with no history of antibiotic use in 10 years preceding stool collection; hCTRL pool - pooled hCTRL donor stool; hABX - human donors with a history of repeated antibiotic use; hABX pool - pooled hABX donor stool. P-value corresponds to unpaired t-test, * - p < 0.05.

### Human-to-mouse faecal microbiota transplantation (FMT)

Male (n=8) and female (n=8) C57Bl/6J 10-week-old mice, originally obtained from Charles River Laboratory Germany were bred in-house and kept in individually ventilated cages in a pathogen-free environment at 22±1°C under a 12:12-hour light-dark cycle. All mice had *ad libitum* access to autoclaved water and food, and were fed a standard chow diet (#801730, Special Diet Services, UK). For FMT, antibiotic pre-treatment to deplete inherent mouse bacteria was performed as previously described (Staley et al., 2017) with modifications (**Figure 2**). Briefly, all mice were given autoclaved drinking water supplemented with an absorbable antibiotic cocktail (ampicillin (1 mg/ml), cefoperazone (0.5 mg/ml), clindamycin (1 mg/ml)) for five days, followed by a two-day washout period. Next, a non-absorbable antibiotic cocktail (streptomycin (1 mg/ml), neomycin (1 mg/ml), vancomycin (0.5 mg/ml)) was administered for an additional five days, followed by a final two-day washout period.

**Figure 2.**
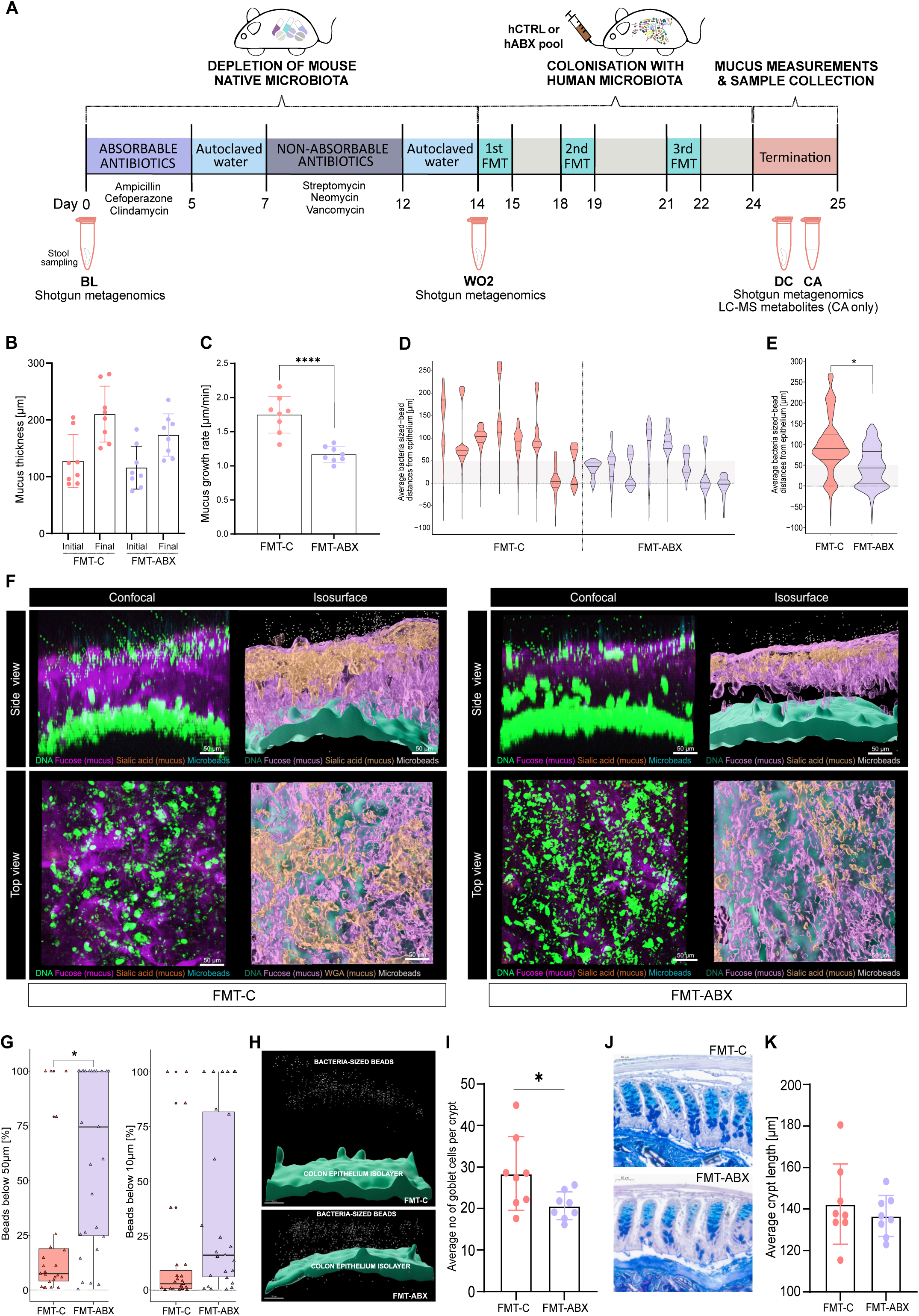
Mucus function analyses of mouse distal colon following FMT. (A) Outline of the mouse experiment; (B) Initial (0 min) and final (45 min) mucus thickness (µm); (C) Mucus growth rate (µm/min, p-value corresponds to unpaired t-test); (D) Average bacteria-sized bead distances from epithelium per mouse (µm); (E) Average bacteria-sized bead distances from epithelium per group (µm, p-value corresponds to linear mixed effects model); (F) Representative confocal Z-stack and isosurface images of the mucosal layer and epithelium in each group; (G) Percentage of beads within 50 µm and 10 µm from the epithelium in each measured image (2-4 images per mouse), per group (p-value corresponds to linear mixed effects model); (H) Representative isosurface images visualising the bead distances from the epithelial isosurface; (I) Average number of goblet cells per crypt (p-value corresponds to Wilcoxon Rank Sum Test); (J) Representative histological images of AB-PAS stained colon tissue to visualize colonic crypts and goblet cells; (K) Average colonic crypt lengths; FMT – Faecal Microbiota Transplant, hCTRL pool – pooled stool from human controls with no history of antibiotic use in 10 years preceding stool collection, hABX pool – pooled stool from human donors with a history of repeated antibiotic use, BL – mouse baseline stool, WO2 – mouse washout 2 stool, DC – distal colon content, CA – caecal content, FMT-C – mice that received FMT from hCTRL pool, FMT-ABX – mice that received FMT from hABX pool.* - p < 0.05; ** - p < 0.01; *** - p < 0.001; **** - p < 0.0001.

Inoculation with human microbiota using oral gavage was carried out directly after the second washout period. FMT gavage suspensions were prepared in an anaerobic chamber (Whitley DG250 Anaerobic Workstation) using 100 mg of human stool sample per 1 ml of PBS supplemented with 0.1% L-cysteine and 15% glycerol (Sigma-Aldrich, St-Louis, MO, USA). The suspension was incubated for 10 min to allow large particles to settle. The four case (hABX) and four control (hCTRL) samples were pooled into two separate suspensions, and 200 µl of suspension per mouse was then gavaged into the corresponding group of antibiotic-pretreated mice (n=4 males and n=4 females per group, n=16 in total). The gavages were repeated after 4 and 7 days. Three days after the final gavage *(i.e.* ten days after the first gavage), the mice were sacrificed, and the distal colon was collected for immediate *ex vivo* analysis of mucus growth rate, mucus penetrability, and histology. Additionally, distal colon content was collected for metagenomic sequencing, caecal content was collected for metabolomics profiling, and mid-colon tissue was collected for RNA expression analyses. Distal colon content was obtained directly from the distal colon upon termination. In case the colon was empty (n = 2), stool samples that were collected one day before the sacrifice were used. Additionally, colon length, caecum weight, abdominal fat weight, and body weight were measured. The researchers were blinded to the case-control groups until the end of the analyses.

### Mucus thickness and growth rate measurements

Colonic mucus layer thickness and growth rate were measured as previously described (Gustafsson et al., 2012; Schroeder et al., 2018). Briefly, the distal part of the colon was gently flushed with Kreb’s buffer (116 mM NaCl, 1.3 mM CaCl2, 3.6 mM KCl, 1.4 mM KH2PO4, 23 mM NaHCO3, and 1.2 mM MgSO4 (pH 7.4)) to remove luminal content and unattached mucus. The muscle layer was then removed, and colonic tissue was mounted in a horizontal perfusion chamber system supplemented with a continuous basolateral supply of RPMI 1640 Medium-Gibco (ThermoFisher Scientific). The surface was visualised by adding black 10 μm polystyrene microspheres (Polysciences, Warrington, USA) on top of the mucus. Kreb’s-mannitol (10 mM mannitol, 5.7 mM sodium pyruvate and 5.1 mM sodium glutamate) was added apically for hydration. Mucus thickness was measured using a fine glass micropipette connected to a micrometre under a stereomicroscope (Olympus). The mucus growth rate (μm/min) was obtained by measuring mucus thickness at time points 0 and 45 min, and then calculating the change in mucus thickness per minute.

### Mucus penetrability measurements

Mucus penetrability was measured as previously described (Gustafsson et al., 2012; Schroeder et al., 2018) with modifications. Briefly, the distal colon tissue was prepared in the same way as described above for mucus growth measurements. After placing the tissue in the perfusion chamber, the epithelium was stained with Syto 9 (ThermoFisher Scientific, 1:500 in Kreb’s-mannitol buffer), and the mucus layer was stained with Wheat Germ Agglutinin (ThermoFisher Scientific, 1:20 in Kreb’s mannitol buffer) and Ulex Europaeus Agglutinin I (Vector Laboratories, 1:20 in Kreb’s-mannitol buffer) for 10 min in the dark, on ice.

Subsequently, the tissue was washed with Kreb’s-mannitol buffer and 1 µm fluorescent microspheres (ThermoFisher Scientific, 1:20 in Kreb’s-mannitol buffer) were added on top of the mucus. The tissue was incubated for a further 10 min in the dark, on ice, to allow time for the microspheres to settle onto the mucus layer. Excess microspheres were gently washed away with Kreb’s-mannitol buffer and the tissue was then covered apically with fresh Kreb’s-mannitol buffer. The tissue was visualised by acquiring z-stack images (5 µm steps) with a 20x/0.5 N-Achroplan water dipping objective on an upright Zeiss LSM 800 confocal microscope. Two to four z-stack images per mouse were taken. The images were then exported and processed with Imaris (Version 9.9.0, Oxford Instruments) to map epithelium, mucus, and microspheres to isosurfaces. On average 916 ± 185 (min 501, max 1252) distances between individual microspheres and epithelial surface per image were extracted and mucus penetrability was quantified by analysis of microsphere distribution within the mucus layer. The average fraction of microspheres penetrating the areas within 10 µm and 50 µm distance to the colonic epithelium were plotted for each image from the mice.

### Tissue histology

From each mouse, a section of distal colon was fixed in Methacarn solution (60% methanol, 30% chloroform, and 10% glacial acetic acid) for at least one week. Samples were then Paraffin-embedded using Sakura Tissue Tek VIP (USA). For Alcian-Blue-Periodic acid-Schiff (AB-PAS) staining, 5 µm colon sections were prepared. In brief, the staining procedure included deparaffinisation of the sections in xylene (VWR Chemicals) and rehydration in ethanol gradients (99%, 90%, and 70%) and water. Thereafter, the slides were placed in 3% acetic acid (VWR Chemicals) and then stained with Alcian-Blue (Sigma-Aldrich) for 20 min. Tissues were oxidised in 0.05% periodic acid (Sigma-Aldrich) before staining with Schiff’s reagent (Sigma-Aldrich) for 20 min. Mayer’s haematoxylin (Sigma-Aldrich) was used for nuclear visualisation, and section dehydration was performed in water, ethanol steps (70%, 90%, and 99%), and xylene. The resulting stained tissues were mounted under coverslips using Pertex glue (Histolab) and images were captured with 20X magnification using Pannoramic Scan P250 Flash III BL/FL (3DHistech, Hungary). The number of goblet cells per crypt and crypt length were evaluated in at least 10 crypts per mouse by three blinded scientists.

### Colony-forming unit (CFU) counts

Stool samples were taken from mice at multiple time points for CFU counting: at baseline before the start of the experiment (Day 0), after the absorbable antibiotic treatment (Day 5), after the first washout period (Day 7), after the non-absorbable antibiotic treatment (Day 12), and after the second washout period (Day 14). Stool samples were weighed and mixed with 500 µl of sterile PBS. The samples were then plated on Brain-heart infusion (BHI) agar plates and incubated under anaerobic conditions at 37°C for 2 days. Thereafter, the colonies were counted.

### RNA extraction and cDNA generation

A biopsy of the distal colon was collected after sacrifice, immediately snap-frozen in liquid nitrogen and stored at -80⁰C until extraction. For RNA extraction, the tissue was homogenised in a TissueLyser II (Qiagen, Germany) using stainless steel beads (5 mm; Qiagen). RNA was then extracted using a RNeasy Mini kit (Qiagen), following the manufacturer’s protocol. RNA concentration and quality were determined using a Nanodrop Lite Spectrophotometer (ThermoFisher Scientific). Per sample, 500 ng of RNA was reverse transcribed to cDNA with a High-Capacity cDNA Reverse Transcription Kit (ThermoFisher Scientific) and diluted 1:7 in nuclease-free water.

### Quantitative real-time PCR analysis

Mouse cDNA was amplified with gene-specific primers and HotStarTaq Master Mix Kit (Qiagen) (Table 1). Amplicons were cloned into a pGEM-T vector (Promega, WI) and transformed into competent DH5α *E. coli* cells. The plasmids were isolated using a Plasmid Mini Kit (Qiagen)*, s*equenced using Sanger sequencing (Eurofins Genomics, Ebersberg, Germany), the target copy number/ng DNA quantified, and target-specific dilution series were prepared. The copy number of specific transcripts was determined by analysing mouse cDNA in a 10 μl reaction mix consisting of 1x iQ SYBR® Green Supermix (Bio-Rad, USA), 0.2 μM of each primer and 2 μl of template cDNA on a CFX Connect Real-Time System (Bio-Rad). Samples and plasmid standards were amplified using the following protocol: denaturation at 95°C for 3 min, followed by 35 cycles of denaturation at 95°C for 20 sec, gene-specific annealing temperature (see Table 1) for 40 sec and extension at 72°C for 1 min. A standard curve was generated for each gene of interest and the transcript copy number in each sample was calculated using the Bio-Rad CFX Maestro software (v.2.3) and reported as copy number/10 ng RNA.

**Table 1.**
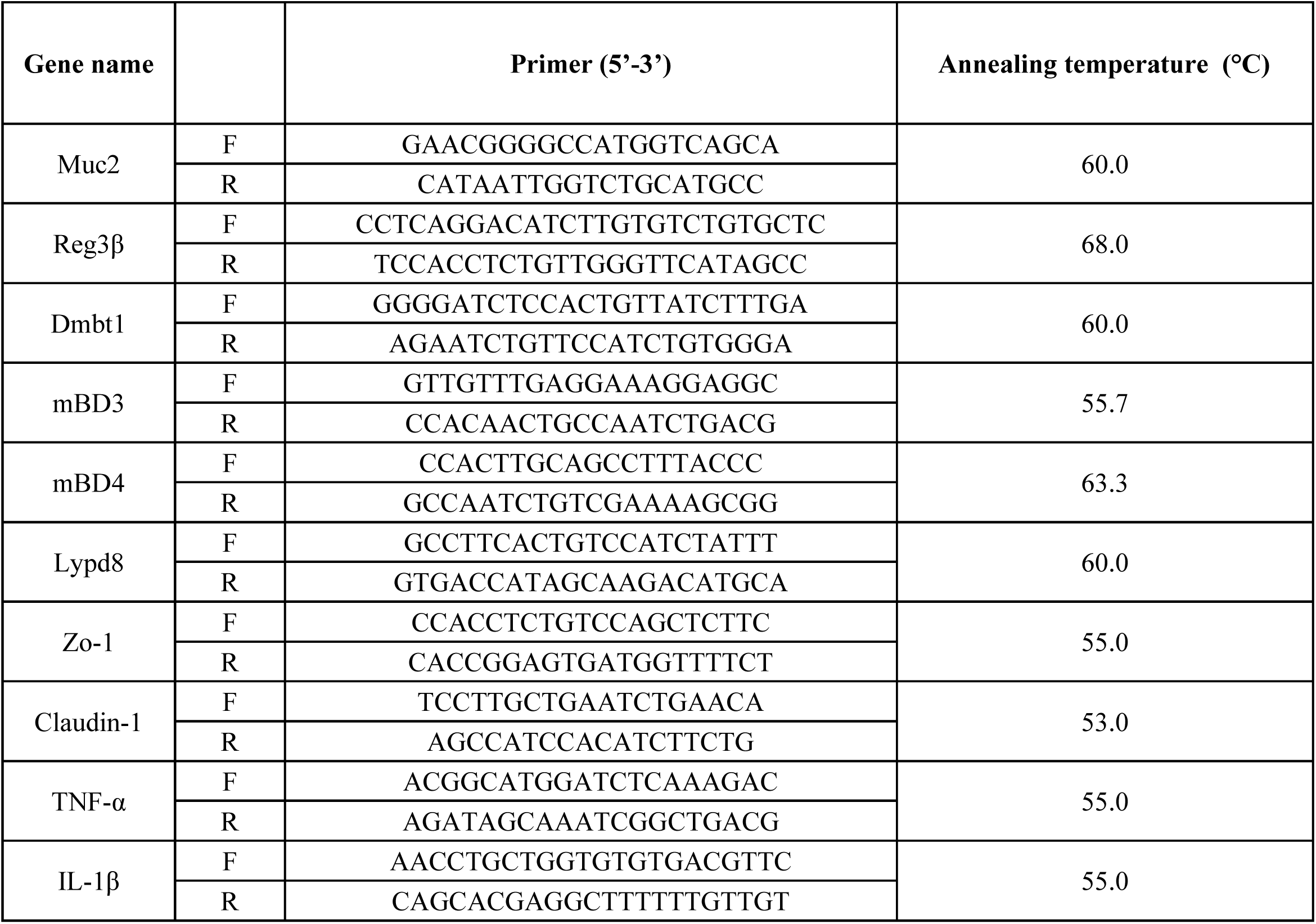
Gene-specific primers and annealing temperatures used for quantitative real-time PCR analysis.

### DNA extraction and metagenomic sequencing

DNA extraction and sequencing of the EstMB donor stool samples was carried out as described previously (Aasmets et al., 2022). For the mouse samples as well as the human stool gavage samples, the microbial DNA extraction was performed using a DNeasy PowerSoil Pro Kit (Qiagen, Germany), according to the manufacturer’s instructions. DNA concentration was measured using the Qubit® dsDNA Assay Kit in Qubit® 2.0 Fluorometer (Life Technologies, CA, USA). The DNA quality control, library preparation, and shotgun metagenomic paired-end sequencing were performed by Novogene Bioinformatics Technology Co., Ltd. Samples were sequenced using the Illumina NovaSeq6000 platform alongside one mock community sample and a negative control.

### Bioinformatics analysis and processing of the metagenomic sequencing data

Reads were trimmed for quality and adapter sequences using fastp (Chen et al., 2018) with - -length_required 5 and --cut_mean_quality 30. The host reads that aligned to the human (build GRCh38) and mouse (build GRCm39) genomes were removed using Bowtie 2 (Langmead and Salzberg, 2012)with parameters --minins 200 and --maxins 400 and SAMtools (Li et al., 2009). This resulted in a total of 763,608,660 paired reads from all mouse samples, excluding controls (n=64, mean 11,931,385 ± 7,366,927 read length 2 x 150 bp) and 116,713,904 paired reads from human donor samples (n=8, mean 14,589,238 ± 1,153,486, read length 2 x 150 bp). Additionally, 14,404,157 reads were obtained from the hCTRL pool sample, and 26,510,699 reads were obtained from the hABX pool sample. As expected, the read numbers after the second washout (n=16, mean 219,232 ± 254,232) were exceptionally low compared to the rest of the samples (n = 61, mean 15,884,751 ± 3,294,077), leading us to exclude this data from further analysis. We did not rarefy the counts to avoid loss of data. The taxonomic composition of the decontaminated and quality-filtered metagenomes was identified by using MetaPhlAn 4 (Blanco-Míguez et al., 2023) with default parameters and database version mpa_vOct22_CHOCOPhlAnSGB_202212. In total, 20 phyla, 858 genera and 1277 species were identified from the sample set.

Additionally, the host-cleaned reads from the human donors and distal colon content samples were also used for *de novo* metagenomic assembly to detect *Akkermansia muciniphila* strains from these samples. First, the reads were assembled into contigs with MEGAHIT v1.2.9 (Li et al., 2015). Thereafter, the contigs were binned using single binners - MetaBAT v2.15 (Kang et al., 2019), Maxbin v2.2.7 (Wu et al., 2016) and VAMB v3.0.7 (Nissen et al., 2021). Since different binning tools reconstruct genomes at various levels of completeness, a bin aggregation software, i.e., DAS Tool v1.1.4 (Sieber et al., 2018), was used to integrate the results of bin predictions made by VAMB, MetaBAT2, and MaxBin2 to optimise the selection on nonredundant, high-quality bin sets using default parameters. Bin quality, including completeness and contamination were estimated using CheckM v2 (Chklovski et al., 2023). Bin statistics, including total size, number of contigs, N50, GC content, etc., were obtained using seqkit v2.3.1 (Shen et al., 2016). Finally, bins were taxonomically annotated using GTDB-Tk v2.3.0, GTDB release number 214 (Chaumeil et al., 2022). The taxonomic position of the assembled genome was determined using GTDB-Tk. All assembled genomes belonging to the *Akkermansia muciniphila* species were clustered at strain level with an average nucleotide identity (ANI) threshold of 99%, as calculated with FastANI v2.09 (Jain et al., 2018). The clustering procedure resulted in two strain-level clusters. The best quality genome from each strain cluster was selected as its representative based on genome completeness, minimal contamination, strain heterogeneity, and assembly N50. The reads from distal colon content and human samples were then mapped against the representative genomes from each cluster.

### Metabolomics analysis

Metabolomics profiling was carried out at the FIMM Metabolomics Unit at the University of Helsinki, Finland and the scientists were blinded to the sample groups. Metabolites were extracted from 30 mg of caecum samples with 400 µL of cold extraction solvent (Acetonitrile: Methanol: Milli-Q Water; 40:40:20, ThermoFisher Scientific). Subsequently, samples were homogenised with three cycles of 30 sec at 5000 rpm at 4°C, centrifuged and supernatants were passed through a Phenomenex Phree Phospholipid removal 96 well plate using robotic vacuum. Sample filtrates were transferred into evaporation tubes and dried under a gentle stream of nitrogen. The dried samples were then re-suspended with 50 µL of cold extraction solvent (Acetonitrile: Methanol: Milli-Q Water; 40:40:20) and analysed for targeted relative profiling and for relative SCFA abundance on a Thermo Vanquish UHPLC coupled with Q-Exactive Orbitrap quadrupole mass spectrometer, equipped with a heated electrospray ionisation (H-ESI) source probe (ThermoFisher Scientific).

Scanning was performed using full MS and polarity switching mode in the mass range 55 to 825 m⁄z and the following settings: resolution of 70,000, spray voltages: 4250 V for positive and 3250 V for negative mode, sheath gas: 25 arbitrary units (AU), and auxiliary gas: 15 AU, sweep gas flow 0, Capillary temperature: 275°C, S-lens RF level: 50.0. Instrument control was operated with Xcalibur 4.1.31.9 software (ThermoFisher Scientific).

The chromatographic separation for metabolite profiling was performed using a SeQuant ZIC-pHILIC (2.1×100 mm, 5-μm particle) column (Merck) at 40°C, with a flow rate of 100 µl/min and a total run time of 24 min. Gradient was started with 2 min at 80% mobile phase B (Acetonitrile), and then gradually increased to 80% mobile phase A (20mM ammonium hydrogen carbonate in water, adjusted to pH 9.4) until 17 min, back to 20% A at 17.1 min and then equilibrated to the initial conditions for 7 min.

For SCFA profiling the separation was performed using a Hypercarb Porous Graphitic Carbon HPLC Column, 50x2.1 3μm (ThermoFisher Scientific) with the gradient starting at 2.5 min at 100% mobile phase A, gradually increased to 100% mobile phase B for 10 min, held until 18 min, returned to 100% A at 18.1 min and then equilibrated to the initial conditions for 6 min. TraceFinder 4.1 software (ThermoFisher Scientific) was used for data integration, the final peak integration and peak area calculation of each metabolite. Data quality was monitored throughout the run using pooled sample as Quality Control (QC), prepared by pooling each study sample, which was interspersed after every 10th sample throughout the run.

Out of the targeted metabolomics panel with 462 metabolites, 220 metabolites were detected in our samples. The metabolite data was checked for peak quality (poor chromatograph), RSD (%relative standard deviation, 20% cutoff) and carryover (20% cutoff). Additionally, metabolites with multiple missing values were excluded from analysis. After quality control, 171 metabolites remained for further analysis. No samples were excluded from the data analysis. These identified metabolites were normalised according to sample weight before analysing the data (peak intensities) using Metaboanalyst 6.0 (www.metaboanalyst.ca).

The metabolomics profile data were transformed using log-transformation. Data scaling was carried out for each variable by mean-centring the variable divided by the square root of the standard deviation (i.e. auto-scaling). Significantly different metabolites between the two groups were identified using the Wilcoxon Rank Sum test. Metabolites with a fold change of 1.5 and an FDR < 0.05 were considered significant.

The same quality procedures were applied to the SCFA metabolomics panel, and all 4 metabolites passed quality control. The metabolites were also normalised according to sample weight before analysing peak intensities, and significantly different metabolites between the two groups were identified using the Wilcoxon Rank Sum test.

### Statistical analyses

If not stated otherwise, data analysis was completed using GraphPad Prism 8. For comparisons between the groups, an unpaired t-test was used when samples were distributed normally, as indicated by the Shapiro-Wilk test, and Mann-Whitney U / Wilcoxon Rank Sum Test was used for non-normally distributed samples. In the case of assessing sex differences in body characteristics between the FMT-ABX and FMT-C groups, comparisons were carried out with multiple t-tests. For correlation analyses, Pearson correlation coefficients were computed for normally distributed data, as determined by the D’Agostino & Pearson test, while Spearman correlation coefficients were computed for non-normally distributed data. P-values were corrected for multiple comparisons where appropriate, according to the Benjamini-Hochberg procedure (False Discovery rate (FDR) < 0.05). In all figures, data are presented as mean ± SD.

Additional analyses were performed using different packages in the R environment (version 4.3.1), as described in further detail below. In all R analyses, stringr (version 1.5.0) and tidyverse (version 2.0) packages were used for data manipulations and ggplot2 (version 3.3.6), ggsci (version 2.9), and ggpubr (version 0.4.0) packages were used for data visualisations.

To evaluate whether there are significant differences in mucus penetrability between the two groups, the linear mixed effects model was used by applying *lmer* function in the lmerTest package (version 3.1.3) on bead-distance measurement values, with the study group as a fixed parameter and mouse and image number as nested random parameters. This allowed us to account for the dependencies in data arising from different numbers of observations per mouse and image.

For the microbiome analyses, the R package phyloseq (version 1.46.0) was used to import, store, and analyse the data. The observed number of unique species (richness) and the Shannon diversity index were used to assess the alpha diversity using the vegan package (v2.6.4). The Euclidean distance on the centred log ratio (CLR)-transformed microbiome species profile was used to calculate the between-sample distances for the beta diversity analysis. Permutational analysis of variance (PERMANOVA) on the Euclidean distances was carried out using the *adonis* function in the vegan package to test the associations between the groups and microbiome composition using 10,000 permutations for the p-value calculations. Alpha and beta diversity analyses were performed on the whole identified composition. To test the differential abundance of different species, Analysis of Compositions of Microbiomes with Bias Correction (ANCOM-BC, package version 2.4.0) and ANOVA-Like Differential Expression (ALDEx, package ALDEx2 version 1.34.0) were used. To limit the number of tests carried out when studying differentially abundant species, the species detected in at least two samples, per sample type, with mean relative abundance higher than 0.1% were selected, resulting in 105 species being tested in distal colon content samples. To account for multiple tests the Benjamini–Hochberg procedure was applied (FDR < 0.05).

## RESULTS

To investigate the potential effect of gut microbiota changes caused by long-term antibiotic use (Aasmets et al., 2022), we carefully selected human subjects from the EstMB cohort using extensive Electronic Health Records (EHR), self-reported microbiome questionnaire data and stringent exclusion criteria (**Fig 1A**, **Supplementary Tables 1 & 2).** Briefly, from 2509 Estonian Microbiome Project participants, four subjects who had not used antibiotics in the 6 months prior to stool sample collection but did use at least five courses of antibiotics in the last five years (hABX), and four subjects with no antibiotics usage history during the last 10 years (hCTRL) were selected. The hCTRL donors were matched with hABX donors based on BMI, sex, age, stool type (Bristol stool scale) and diet, as these factors are known to be major drivers of microbiome variability (**Table 2**). All selected participants were healthy, with normal BMI and no history of complex diseases (41 excluded diseases, **Supplementary Table 1**) nor any history of recent medication usage (27 excluded medications, **Supplementary Table 2**) that had been previously associated with the microbiome composition (Aasmets et al., 2022).

**Table 2.**
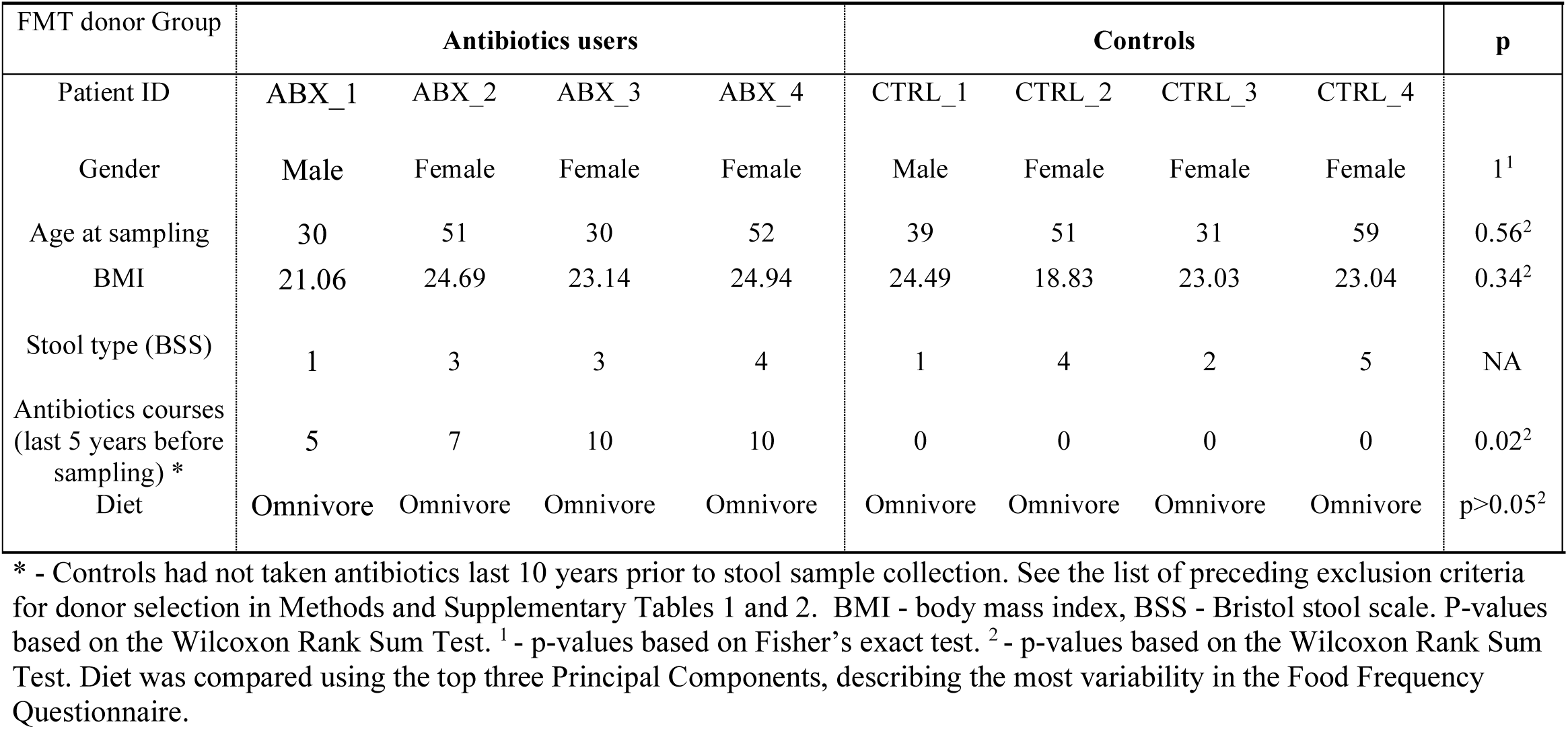
Characteristics of the Faecal Microbiota Transplant donors.

| FMT donor Group | Antibiotics users |  |  |  | Controls |  |  |  | p |
| --- | --- | --- | --- | --- | --- | --- | --- | --- | --- |
| Patient ID | ABX_1 | ABX_2 | ABX_3 | ABX_4 | CTRL_1 | CTRL_2 | CTRL_3 | CTRL_4 |  |
| Gender | Male | Female | Female | Female | Male | Female | Female | Female | 1<sup>1</sup> |
| Age at sampling | 30 | 51 | 30 | 52 | 39 | 51 | 31 | 59 | 0.56<sup>2</sup> |
| BMI | 21.06 | 24.69 | 23.14 | 24.94 | 24.49 | 18.83 | 23.03 | 23.04 | 0.34<sup>2</sup> |
| Stool type (BSS) | 1 | 3 | 3 | 4 | 1 | 4 | 2 | 5 | NA |
| Antibiotics courses<br>(last 5 years before<br>sampling) * | 5 | 7 | 10 | 10 | 0 | 0 | 0 | 0 | 0.02<sup>2</sup> |
| Diet | Omnivore | Omnivore | Omnivore | Omnivore | Omnivore | Omnivore | Omnivore | Omnivore | p>0.05<sup>2</sup> |

\* - Controls had not taken antibiotics last 10 years prior to stool sample collection. See the list of preceding exclusion criteria for donor selection in Methods and Supplementary Tables 1 and 2. BMI - body mass index, BSS - Bristol stool scale. P-values based on the Wilcoxon Rank Sum Test. <sup>1</sup> - p-values based on Fisher's exact test. <sup>2</sup> - p-values based on the Wilcoxon Rank Sum Test. Diet was compared using the top three Principal Components, describing the most variability in the Food Frequency Questionnaire.

Interestingly, although both donor groups were generally healthy and did not use antibiotics for at least 6 months prior to stool sample collection, analysis of microbial communities identified a significantly lower number of observed microbial species in the hABX donors compared to the hCTRL donors (168.0 ± 77.69 [hABX] versus 322.2 ± 46.54 [hCTRL]; p = 0.02) (**Fig 1B**). A similar, yet not statistically significant, trend was observed for the Shannon diversity index (2.9 ± 1.13 [hABX] versus 4.1 ± 0.21 [hCTRL]; p = 0.11) (**Fig 1B**). Furthermore, the principal component analysis biplot shows the clustering of these two groups, though this difference did not reach statistical significance according to the permutational analysis of variance (PERMANOVA R^2^ = 0.1856, p = 0.085) (**Fig 1C**). Analysis of gut microbial composition on genus level revealed several significantly different genera between hABX and hCTRL samples, including *Bacteroides*, *Phocaeicola, Parabacteroides, Blautia, Clostridium* and *Akkermansia* among others (p < 0.05, **Fig 1D, Supplementary Table 3**). Importantly, we observed that the donors chosen for this study exhibit similar changes in *Bacteroides* abundance (mean relative abundances 35.27 ± 18 % in hABX donors versus 9.24 ± 7.3% hCTRLs) to those observed across the entire EstMB population cohort, where antibiotic usage resulted in more *Bacteroides*-dominant communities (Aasmets et al., 2022).

### Repeated antibiotic use in humans promotes mucosal barrier dysfunction in mice following FMT

To investigate whether the long-term antibiotic-shaped gut microbiota could contribute to intestinal mucus dysfunction, the stool samples from the hABX and hCTRL participants were pooled in two separate suspensions, according to the respective groups, for faecal microbiota transplantation (FMT). Each FMT group consisted of eight mice (four males and four females), and all mice received the same standard chow diet before and throughout the experiment. Groups of mice receiving microbiota from hABX and hCTRL donors were named FMT-ABX and FMT-C mice, respectively (**Fig 1A**). FMT was performed in microbiota-depleted young adult mice as, unlike germ-free mice, they have undergone normal development of the mucus layer and do not have an under-stimulated intestinal immune system (Johansson et al., 2015; Thompson and Trexler, 1971). Following three repeats of the FMT, colonic mucus function was analysed *ex vivo* on viable tissue 10 days after the first human-to-mouse transplant **(Fig 2A**).

Mucus growth is an intrinsic part of mucus function, which helps to maintain the mucus layer by providing a constant flow from the epithelial surface towards the lumen, thereby pushing microbes and debris away from the proximity of the epithelium. Thus, to compare the mucus growth between the two microbiota-transplanted groups, we measured mucus thickness repeatedly over 45 min to calculate the mucus growth rate. The initial mucus thickness did not differ between the two groups after gently flushing away loose luminal material (116 µm ± 38 [FMT-ABX] versus 128 µm ± 47 [FMT-C]; p = 0.5824), but the thickness after 45 min increased to 173 µm ± 37 in the FMT-ABX group and 210 µm ± 49 in the FMT-C group (p = 0.1142) (**Fig 2B**). Accordingly, the mucus growth rate was significantly lower in the distal colon of hABX mice that received FMT from donors with antibiotic use history (hABX), compared to FMT-C mice that received FMT from healthy controls (hCTRL) (1.16 µm/min ± 0.11 [FMT-ABX] versus 1.74 µm/min ± 0.27 [FMT-C]; p < 0.0001) (**Fig 2C**), allowing the possibility for the microbes to move closer to the epithelium as the mucus flow is not pushing the microbes away as effectively.

While a healthy colonic mucus layer is impenetrable to bacteria, microbes can gain access to the colonic epithelium through degradation and penetration of the glycan-rich gel matrix. To assess mucus penetrability, we collected confocal Z-stack images of mouse distal colon tissue to determine mucus structure and location of bacteria-sized microspheres (i.e. beads) in the mucus. Physical changes to the mucus layer were observed in the top and the side view of the confocal microscopy Z-stack images, with a sparser mucus layer in the FMT-ABX compared to the FMT-C group (**Fig 2F**). Likewise, when measuring the distances of individual beads to the colonic epithelium surface, most of the beads were situated closer to the epithelium in FMT-ABX mice compared to the FMT-C group (**Fig 2D**). Further, the average distance of the beads from the epithelium was significantly lower (p = 0.0246) in the FMT-ABX mice (47±47 µm; median 45 µm) when compared to the FMT-C mice (98±68 µm; median 90 µm) (**Fig 2E**).

In healthy mice, the inner ∼50 µm of the mucus is generally considered impenetrable to microbes (Johansson et al., 2008). However, we observed that an average of 61 ± 39% beads (median 74.55%) were closer than 50 µm from the epithelial surface in the FMT-ABX group, while in the FMT-C group only an average of 23 ± 34% (median 7.51%) were found to be in this vicinity (p=0.0383, **Fig 2G**). These findings suggest an increase in mucus penetrability to microbes in the mouse gut, when colonised by microbiota from humans repeatedly exposed to antibiotics. A similar tendency was observed for the area within 10 µm from the epithelial surface (average 35 ± 40% beads (median 16%) in the FMT-ABX group vs average 17 ± 32% (median 3%), in the FMT-C group), though this difference in proportions of beads did not reach statistical significance (p = 0.22, **Fig 2G**). Correspondingly, visualising the epithelial isosurface and the beads illustrated that the beads were indeed more spread throughout the mucus layer in the FMT-ABX group compared to the FMT-C group, where beads were instead located at a greater distance from the epithelial isosurface **(Fig 2H)**.

Furthermore, counting of mucus-producing goblet cells indicated that the FMT-ABX group had a lower number of filled goblet cells compared to the FMT-C (p = 0.035, **Fig 2I**). Yet, analysis of histological sections did not reveal any signs of inflammation in either group (**Fig 2J**), which was also supported by similar lengths of the colonic crypts (p = 0.57. **Fig 2K**). Overall, these results indicate a change in the physical structure of the mucus layer in the FMT-ABX group, allowing microbes to move closer to the epithelial layer compared to the FMT-C group, supported by both *ex vivo* mucus analyses and histological analysis.

### Increased expression of Muc2 and Reg3γ might compensate for impaired mucus function

As mucus function was impaired in the FMT-ABX compared to the FMT-C mice, we thus wondered whether the deterioration in host defence led to compensatory activation of alternative intestinal defence mechanisms to prevent the bacteria from moving closer to the epithelium. The host can respond to bacterial proximity to the colonic epithelial by increasing the secretion of antimicrobial peptides (AMPs), including defensins, lysozyme, regenerating islet-derived protein 3 gamma (Reg3γ) and Ly6/PLAUR domain containing 8 (Lypd8) (Okumura et al., 2016; Salzman, 2010; Vaishnava et al., 2011). As such, we assessed the expression of Muc2 and these AMPs using absolute quantification of RNA from colonic tissue samples. Expression of Mucin 2 (Muc2), the main component of mucus, was significantly higher in the FMT-ABX group compared to the FMT-C group (p = 0.02, **Fig 3A**). In addition, the expression level of the antibacterial lectin Reg3γ was significantly higher in the FMT-ABX group (mean 1.6×10^3^ transcripts/10ng RNA [FMT-ABX] versus 4.1×10^2^ transcripts/10ng RNA [FMT-C]; p=0.003, **Fig 3B**). However, expression of additional intestinal mucosal host defence proteins, such as lysozyme (Lys), mouse beta-defensin 4 (mBD4), Lypd8 or Deleted in malignant brain tumours 1 (Dmbt1) was not significantly different between the two groups **(Fig 3C-F).** Likewise, expression of the tight junction proteins Zonulin-1 and Claudin-1, as well as inflammatory markers TNF-α and IL-1β was also not significantly different between the two groups **(Supplementary Fig 1).** Consequently, the microbiota-mediated mucus defect led to a selective compensatory host response towards encroaching gut bacteria. Moreover, linking the expression data of Muc2 with our mucus function measurements demonstrates that measuring Muc2 expression without any functional mucus analysis can be misleading, as it disregards the relevant post-transcriptional processes of mucus proteins that influence mucus properties and function.

**Figure 3.**
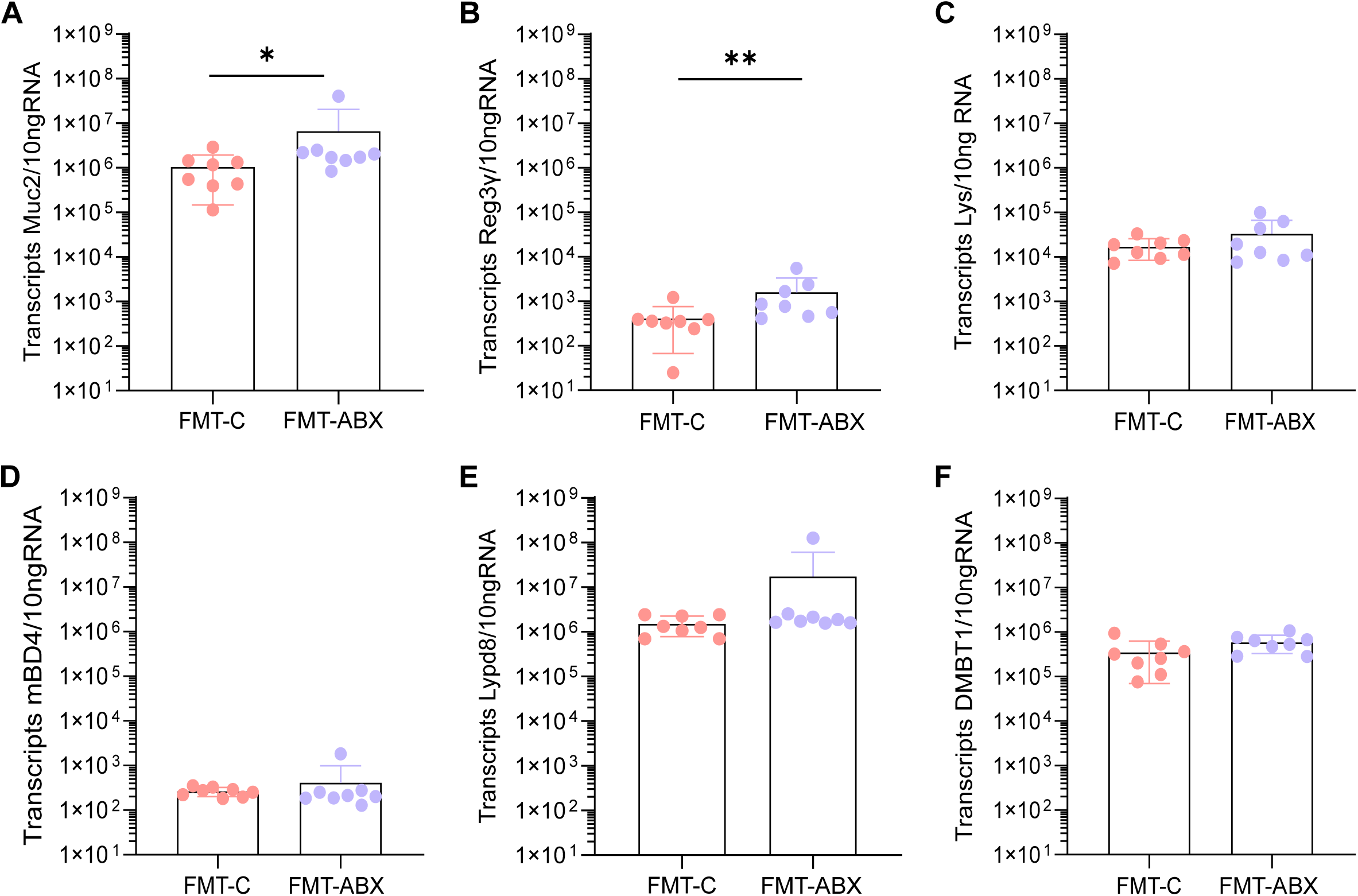
Absolute RNA expression of Muc2 and antimicrobial peptides in mouse mid-colon following FMT. Absolute quantification of RNA expression levels of (A) Mucin 2 (Muc2); (B) Regenerating islet-derived protein 3-gamma (Reg3γ); (C) Lysozyme (Lys); (D) Mouse Beta-defensin 4 (mBD4); (E) LY6/PLAUR domain containing 8 (Lypd8); (F) Deleted in malignant brain tumours 1 (Dmbt1) in mouse mid-colon tissue; FMT – Faecal Microbiota Transplant. All p-values correspond to the Wilcoxon Rank Sum Test. * - p < 0.05; ** - p < 0.01.

### Physiological parameters and distal colon microbiome composition differ between antibiotic users and healthy controls

Antibiotic-mediated changes in the gut microbiota are linked to metabolic impairments in humans and mice (Fenneman et al., 2023). To characterise whether the transplanted microbiome shaped by a history of antibiotic usage affected the mouse metabolism, we assessed the weight of abdominal fat and total body weight. Strikingly, we observed a significantly higher weight of abdominal fat in FMT-ABX mice (p = 0.028, **Fig 4A**), despite total body weight not differing between the two groups (**Fig 4B**). Furthermore, the percentage of abdominal fat contributing to total body weight was significantly higher in FMT-ABX mice (p = 0.037, **Fig 4C**).

**Figure 4.**
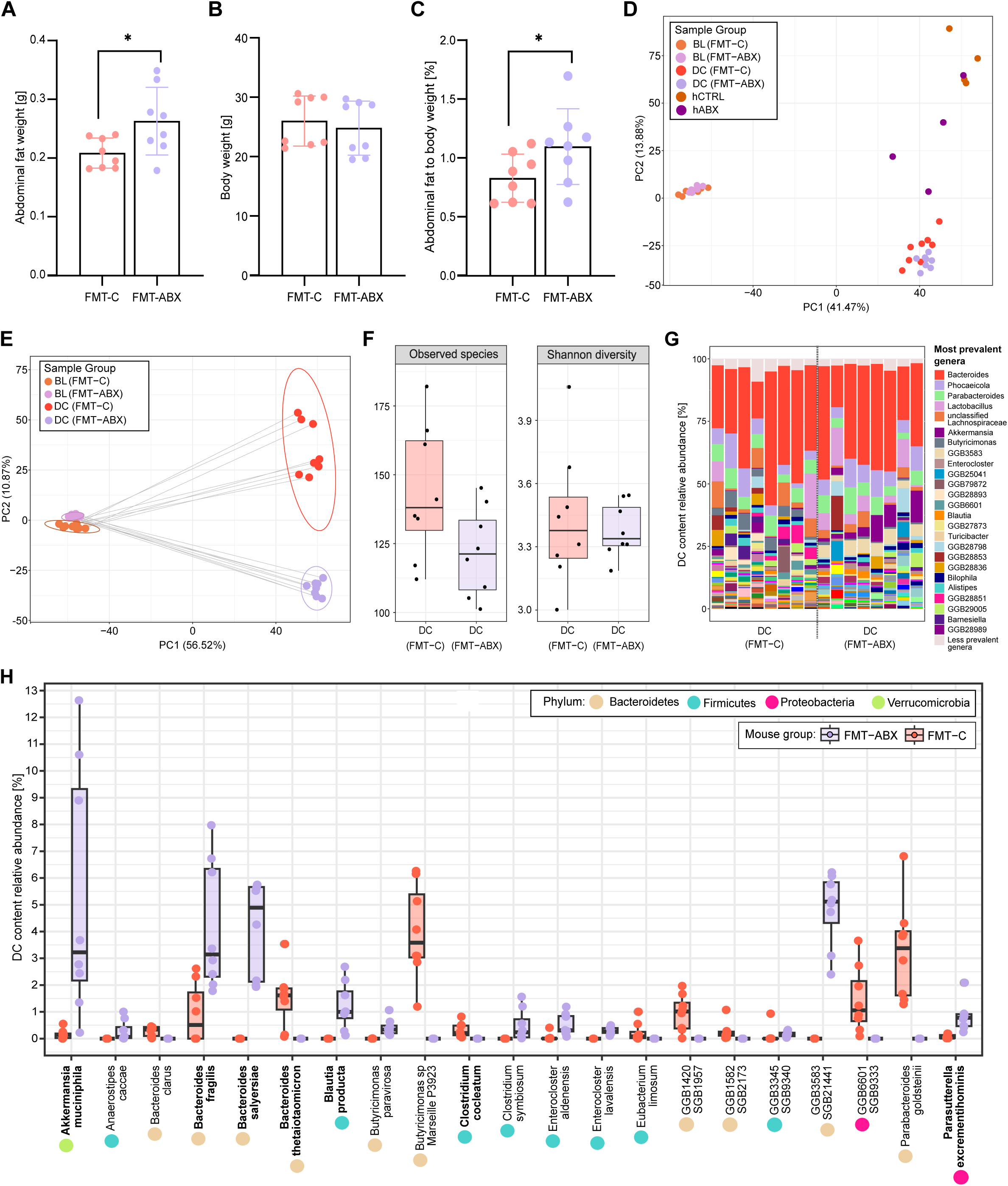
Physiological parameters and distal colon microbiome composition of mice following FMT. (A) Abdominal fat weight (unpaired t-test) and (B) total body weight after FMT; (C) Abdominal fat as a percentage of total body weight (Wilcoxon Rank Sum Test); (D) Beta diversity of the microbiota between FMT-C and FMT-ABX mouse stool at baseline, DC content after FMT and human donor stool used for FMT; (E) Beta diversity of the microbiota between FMT-C and FMT-ABX mouse stool at baseline and DC content after FMT; (F) Boxplots representing alpha diversity (observed species and Shannon diversity index) of the mouse DC content; (G) Most prevalent genera in FMT-C and FMT-ABX DC content of individual mouse samples; (H) Relative abundances of significant differentially abundant species in mouse DC content between the two groups (FDR < 0.05). Bacterial species previously shown to degrade mucus are indicated in bold. FMT – Faecal Microbiota Transplant. BL - mouse baseline stool, DC – mouse distal colon content, hCTRL - human controls with no antibiotic use history in last 10 years, hABX - human donors with a history of repeated antibiotic use, FMT-ABX - mice receiving FMT from hABX pool, FMT-C - mice receiving FMT from hCTRL pool. * - p < 0.05.

To next identify the differences in the microbial communities between the FMT-ABX and FMT-C groups, we initially evaluated the success of the microbiota transplant. Depletion of the inherent mouse microbiota prior to FMT by antibiotic cocktails was confirmed by stool plating, displaying a reduction in colony-forming units (CFU/g of stool) by at least 6 logs (**Supplementary Fig 2**). Additionally, through plating of the pooled samples/FMT material, we confirmed that the microbes in the pooled donor samples were viable in both groups (> 10^8^ CFU/g of pooled stool). Moreover, shotgun metagenomic sequencing identified that the mouse microbiota at baseline (BL, Day 0), i.e. before microbiota depletion, was significantly different from human donor samples (PERMANOVA R^2^ = 0.5319, p < 0.00005), as well as from distal colon (DC) content taken from the mice at termination, 10 days after first colonisation with human microbiota (PERMANOVA R^2^ =0.56335, p < 0.00005). Correspondingly, the baseline mouse microbiota separated from the human donors and mouse DC content samples as shown by the first principal component, indicating a replacement of the mouse baseline microbiota by a community that rather resembles the human microbiota (**Fig 4D**). Crucially, Euclidian distance metric-based beta diversity analysis of the CLR-transformed species-level profile confirmed that the FMT transplant led to a significantly different, and donor-group specific, microbiome composition of the mice **(Fig 4E**, PERMANOVA R^2^ = 0.3163, p<0.0002). Similar to the differences seen in the human donor groups, a lower number of species was observed in the FMT-ABX mice compared to the FMT-C mice though the difference was not statistically significant (p = 0.056 and p = 0.68), as was also the case for the Shannon diversity index **(Fig 4F**).

While no striking overall differences between individual FMT-ABX and FMT-C DC content samples were observed for the most common genera **(Fig 4G**), compositional profiling at species level revealed that 21 species from 4 different phyla were differentially abundant between the two groups (FDR < 0.05, **Fig 4H**). Of these 21 species, 12 were more abundant in the FMT-ABX group and 9 in the FMT-C group. In fact, the 9 species were found to be unique to the FMT-C group, having not been detected in the FMT-ABX samples, which may indicate that these species play a protective role against development of a mucus deteriorating phenotype (**Fig 4H**). Conversely, out of the 12 species that were enriched in the FMT-ABX group, 6 could not be detected in any of the FMT-C samples (**Fig 4H**), again indicating that antibiotic usage might induce compositional changes, resulting in communities distinct from those of healthy controls. Interestingly, multiple species that were dominant in the FMT-ABX have been previously shown to be mucin utilisers, including *Akkermansia muciniphila*, *Blautia producta*, *Parasutterella excrementihominis,* and species of the *Bacteroides* genus, such as *Bacteroides fragilis* and *Bacteroides salyersiae* (Glover et al., 2022; Pan et al., 2022). Previously reported mucin-utiliser species i.e. *Bacteroides thetaiotaomicron* (Bry et al., 1996) and *Clostridium cocleatum* (Boureau et al., 1993) were also found to be significantly more abundant in the FMT-C group, however, their relative abundance in the community was much lower, compared with the mucin utilisers in the FMT-ABX group which seemed to dominate in the community.

### History of antibiotic use results in a distinct microbial metabolite profile

The health status of the gut environment is determined by both the gut microbiota and the host immune system, as well as the interactions between them. The gut microbiota signal to the host via the production of metabolites (Schroeder and Bäckhed, 2016). To thus analyse whether the history of antibiotics usage affects the microbial metabolomics profile in the transplanted mice, we carried out relative metabolomics profiling on mouse caecal content using high-performance liquid chromatography–mass spectrometry. Unsupervised hierarchical cluster analysis of the 171 reliably detected metabolites identified a clear separation based on transplant group (**Fig 5A**), with distinct clusters of metabolites being enriched or depleted in either group. 10 metabolites (5.8%) showed significantly different abundance levels between the FMT-ABX and FMT-C groups, with the majority (adenine, adenosine, betaine, butanoate, 4-coumarate, dihydroxybenzoate, 4-imidazole acetic acid, propionate, pentanoate) being enriched in the FMT-ABX group (**Fig 5B**, FDR < 0.05). In contrast, N-Acetyl-L-Alanine was the only metabolite that was significantly enriched in the FMT-C group. Consequently, these findings indicate that a history of antibiotic use can result in significant alterations of microbial metabolism, compared with healthy controls.

**Figure 5.**
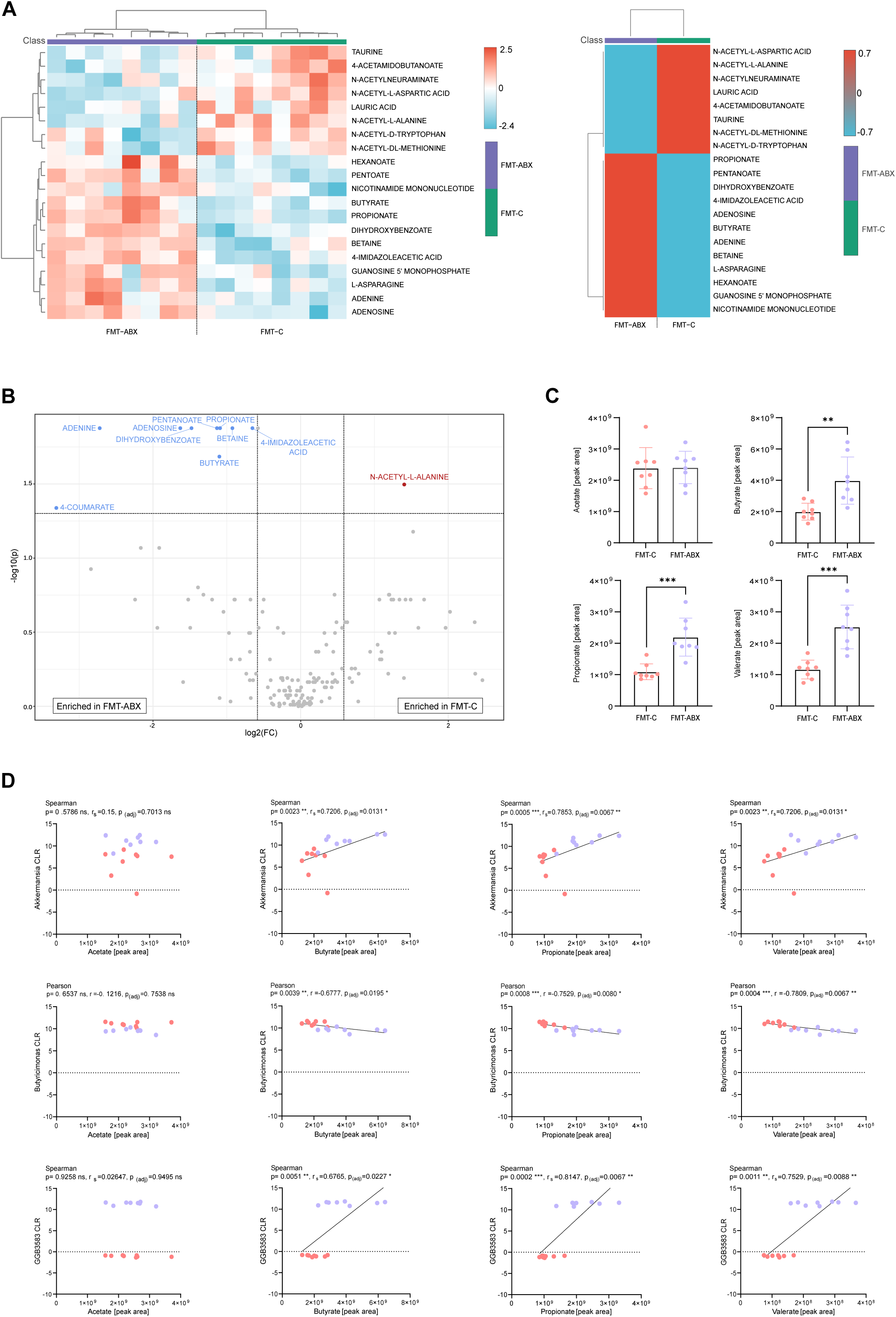
Metabolomic profiling of mice following FMT. (A) Individual mouse and FMT-grouped heatmaps of top 20 metabolites from relative metabolomics profile (unpaired t-test p < 0.05); (B) Significant differentially abundant metabolites in relative metabolomics profile (FDR < 0.05); (C) Targeted SCFA metabolomics analysis of acetic acid, propionic acid, butyric acid, valeric acid, isovaleric acid and 2-methylvaleric acid; (D) Correlation between SCFA peak area and CLR of *Akkermansia*, *Butryicimonas* and *GGB3583*, p-values correspond to Spearman or Pearson correlation, as noted, p-value correction was performed using the Benjamini-Hochberg procedure (FDR < 0.05).; All other p-values correspond to the Wilcoxon Rank Sum Test. * - p < 0.05; ** - p < 0.01, *** - p < 0.001. CLR = Centred Log Ratio.

Among the differentially abundant metabolites, we identified the short-chain fatty acids (SCFAs) propionate, butyrate and pentanoate. As SCFAs have been associated with gut health and mucosal barrier function (Holmberg et al., 2023; Liu et al., 2021; Willemsen et al., 2003) we further quantified the SCFA levels in the different groups by targeted short-chain fatty acid (SCFA) profiling. Confirming the global profiling (**Fig 5A**), propionic, butyric and valeric acid were found to be significantly higher in mice who received the FMT from human donors with a history of repeated antibiotic use (p < 0.05, **Fig 5C**), which was unexpected with regards to the general assumption that higher levels of SCFAs correlate with improved mucosal barrier function. To thus explore this surprising observation further, we tested for potential correlations between the SCFAs and the most abundant microbial genera. After adjusting for multiple comparisons, we found that *Akkermansia, Butyricimonas* and *GGB3583* each significantly correlated with propionate (p(adj)=0.0067, rs=0.7853; p(adj)=0.0080, r=-0.7529; p(adj)=0.0067, rs=0.8147), butyrate (p(adj)=0.0131, rs=0.7206; p(adj)=0.0195, r=-0.6777; p(adj)=0.0227, rs=0.6765) and valerate (p(adj)=0.0131, rs=0.7206; p(adj)=0.0067, r=-0.7809; p(adj)=0.0088, rs=0.7529 ), but not acetate levels, respectively (**Fig 5D**). However, due to the group-dependent differences in CLR abundances for GGB3583, we considered the correlations of this genus as a methodological artefact. Correlations for the remaining most abundant genera were not significant (**Supplementary Fig 3**).

*Akkermansia* correlated positively with propionate, butyrate and valerate production, particularly in the FMT-ABX group where these SCFA concentrations were higher. Interestingly, metagenomics analyses confirmed that the *Akkermansia* genus in our dataset consisted of *Akkermansia muciniphila*, a well-known mucin degrader (Derrien et al., 2004), thereby corroborating previous findings that *A. muciniphila* mucolytic activity is associated with the production of SCFAs (Derrien et al., 2004; Ottman et al., 2017). However, when testing for correlations within the two individual groups we detected correlations in opposite directions in the FMT-ABX group (propionate p=0.003, r=0.7561; butyrate: p=0.0495, r=0.7078; valerate: p=0.5507, r=0.2498) when compared to the control FMT-C group (propionate p=0.793, rs=-0.119; butyrate: p=0.6191, rs=-0.2143; valerate: p=0.793, rs=0.119). Intrigued by this observation, we performed *de novo* metagenomic assembly and identified two distantly related *A. muciniphila* strains, indicated by an average nucleotide identity (ANI) index >99%, that were distinct between the mice receiving FMT from donors with a history of antibiotics users and respective controls. While further characterisation of these two strains is required, it is thus possible that repeated antibiotics usage selects for a gut microbial community that is detrimental to mucus function.

## DISCUSSION

Maintaining a safe distance between gut microbiota and the epithelial barrier is a crucial characteristic of intestinal health. Recently, impaired mucus barrier function has been connected to different diseases, including ulcerative colitis (Johansson et al., 2014; Van Der Post et al., 2019) and metabolic disease (Chassaing et al., 2017) in humans and has been associated with cancer development in mice (Velcich et al., 2002). Moreover, antibiotic use has been shown to have long-term effects on microbiome composition (Aasmets et al., 2022) and has also been characterised as a risk factor for several diseases such as type-2 diabetes, IBD, and coeliac disease (Sander et al., 2019; Faye et al., 2023; Mamieva et al., 2022; Mikkelsen et al., 2015). However, the link between long-term antibiotic usage and mucus function has so far not been studied.

Here we show that human-derived microbiota from donors with a history of repeated antibiotic use, but no diagnosed disease, can trigger mucus barrier dysfunction in mice, including increased mucus penetrability and reduced mucus growth rate. Mucus growth is a key characteristic of a healthy mucus layer which creates a constant luminal-directed flow of mucus away from the host epithelial layer and replenishes the mucus layer after microbial consumption of the mucin glycans. We observed a significantly lower mucus growth rate in the mice transplanted with microbiota from donors with a history of repeated antibiotic use, compared to controls. This was accompanied by increased penetrability to bacteria-sized beads in the FMT-ABX mice, as evidenced by the proximity of the beads to the epithelium and a higher proportion of beads within the inner area of the mucus, as well as a significantly lower number of filled goblet cells. This indicates several aspects of mucus barrier failure in mice receiving microbiota from donors with a history of antibiotic usage, as a result of both physical and functional alterations.

To counteract the dysfunctional mucus barrier, the host could activate compensatory mechanisms to re-establish protection against the encroaching antibiotics-induced microbial community. In accordance with this, we detected increased mRNA expression of the mucin protein Muc2 and the antimicrobial peptide Reg3γ (the mouse homologue for human REG3A) in the colon of mice transplanted with the microbiota derived from repeated antibiotic users. Expression of both Muc2 and Reg3γ has been previously shown to be induced by the presence of gut bacteria (Vaishnava et al., 2011, 2008), and Reg3γ has also been shown to be highly expressed in the mouse colon during acute colitis (Darnaud et al., 2018), suggesting that the ABX-shaped microbiota may induce a similar host response to that observed in the inflamed gut. Of note, while our observation of a reduced mucus growth rate but increased Muc2 expression in the FMT-ABX mice initially seemed contradictory, this could be explained by the dominance of known mucin-degrading species in this group, with excessive degradation resulting in a lower mucus growth rate despite higher Muc2 expression. Moreover, mucus secretion is regulated post-translationally through autophagy and ER stress (Naama et al., 2023), which is not reflected by measuring Muc2 expression.

Previous studies have highlighted the importance of microbial colonisation for normal mucus barrier function (Johansson et al., 2015) and have implicated certain microbiota and their metabolites in determining Muc2 structure, with conventionally raised mice generally having longer Muc2 glycans than germ-free mice (Arike et al., 2017). Furthermore, microbial composition has also been implicated in mucus layer structure, as demonstrated in studies where the same mouse strain was colonised with different microbiota, resulting in differing mucus function phenotypes (Jakobsson et al., 2015; Rodrıguez-Pineiro and Johansson, 2015). Analysis of the microbiota composition in our transplanted mice revealed that several species present in the FMT-ABX group were absent from the microbial community of the FMT-C control group, suggesting that antibiotics might select for a microbial community that is detrimental to mucus function. Supporting this hypothesis, several species that have previously been shown to degrade mucus differed between the two transplantation groups. Namely, *A. muciniphila* (Derrien et al., 2004), *B. fragilis, B. salyersiae, B. producta*, *and P. excrementihominis* (Glover et al., 2022; Pan et al., 2022), which were more prevalent in the FMT-ABX group when compared to the control group. In contrast, in the FMT-C mice, only *B. thetaiotaomicron* (Bry et al., 1996) and *C. cocleatum* (Boureau et al., 1993) have been previously associated with mucus consumption and, though generally being low in abundance, have higher abundance when compared to the FMT-ABX group.

While moderate mucus consumption is part of a homeostatic microbiota-host interaction, an overly active mucus-foraging microbial community will disturb the balance between mucus secretion and mucus degradation, consequently leading to increased penetrability and barrier breakdown. As such, some bacterial species, including *A. muciniphila*, have been characterised as mucin specialists (Desai et al., 2016), feeding exclusively on mucin O-glycans as a nutrient source, while others have been described as mucin generalists, which are capable of nutrient-dependent flexible foraging. One example of the latter is *B. thetaiotaomicron*, which has a preference for dietary polysaccharides but can switch to feed on mucin o-glycans when dietary nutritional sources are unavailable (Sonnenburg et al., 2005). Additionally, *B. thetaiotaomicron* has been shown to produce SCFAs including acetate and propionate (Wrzosek et al., 2013), and we could recently link these SCFAs to improved mucus growth rate under Western-style diet consumption (Holmberg et al., 2023). Furthermore, *B. thetaiotaomicron* is able to stimulate mucus fucosylation, with induction of fucosylation only occurring if the bacterium is also capable of fucose foraging (Bry et al., 1996). Interestingly, the mucus-promoting abilities of *B. thetaiotaomicron* can be attenuated by the presence of *Faecalibacterium prausnitzii*, demonstrating a homeostatic cross-feeding relationship between bacterial species in the gut in relation to mucus function (Wrzosek et al., 2013).

*A. muciniphila,* which was more abundant in the mouse group that received microbiota from donors with a history of antibiotic use, is a well-known mucus degrader, and its excessive colonisation has shown to break the dynamic balance between mucin production and consumption, thereby affecting intestinal barrier function (Qu et al., 2023). Additionally, previous studies have linked *A. muciniphila*-mediated mucus disruption to an exacerbation of food allergy (Parrish et al., 2023) and *Salmonella typhimurium* infection (Ganesh et al., 2013). In accordance with our findings, the latter study also detected increased Muc2 expression, but lower numbers of mucin-filled goblet cells in the presence of *A. muciniphila*, confirming that Muc2 expression might not be an optimal indicator of mucin production.

*A. muciniphila,* as well as *B. fragilis,* has been shown to repopulate the colon after antibiotic supplementation (Derrien et al., 2017; Lee et al., 2013). In line with this, we detected *A. muciniphila* in the FMT-C mice in modest abundance, while it reached higher abundance in the FMT-ABX group. Interestingly, using *de novo* metagenomic assembly, we could detect two different previously undescribed *A. muciniphila* strains that were distinct between the mice receiving FMT from donors with a history of antibiotics users and respective controls. It is thus possible that strain-specific characteristics led to different establishment within the microbial communities, thereby leading to the distinct physiological phenotypes.

Besides differences in the microbial communities, we detected group-specific metabolomics profiles, characterised by significant differences in the abundance of most of the SCFAs. Unexpectedly, propionate, butyrate and valerate were more abundant in the intestinal content of the FMT-ABX mice, which had a dysfunctional mucus layer. Yet, this may be explained by the fact that most studies that have previously investigated mucus function focused on dietary interventions, often with varying dietary fibre content, which directly affects microbial fermentation and SCFA production (Holmberg et al., 2023; Desai et al., 2016; Schroeder et al., 2018; Riva et al., 2019; Chen et al., 2022). In contrast, when studying mucus function in mouse models of diet-independent obesity, where both groups were fed on a chow diet, no differences in SCFA levels were detected between mouse groups, despite their strong difference in mucus function (Schroeder et al., 2020). Combined with this study, these results suggest that SCFAs can potently modulate mucus function under specific environmental conditions, including defined diet-shaped microbial communities, but that high SCFA levels alone are not sufficient to maintain healthy mucus function. As such, *A. muciniphila* has been shown to produce propionic acid as well as stimulate butyric acid production in syntrophic interactions with *Anaerostipes caccae* (Belzer et al., 2017). Interestingly, in our experiment *A. caccae* was only detected in the mice who received FMT from donors with history of antibiotics use. It is thus possible that the co-occurrence of the two species in an antibiotic-disturbed background community leads to a microbial composition, in which the SCFA-mediated mucus production (Liu et al., 2022) is no longer sufficient to compensate for the excessive mucin degradation by *A. muciniphila.* Alternatively, defects in the intestinal mucosal barrier may impede SCFA-mediated mucus production or SCFA uptake by the intestinal epithelium.

Antibiotic usage early in life is known to increase the risk of developing obesity and central adiposity in humans and mice (Azad et al., 2014; Cho et al., 2012). Interestingly, we observed significantly higher abdominal fat weight in FMT-ABX mice already 10 days after the microbial transplant. At this time point, however, no difference in body weight was observed. Mucus dysfunction has previously been linked to an obesity-associated microbiota in mice, independent of dietary composition (Schroeder et al., 2020), similar to our current study. Moreover, mice deficient in the antibacterial mucus protein zymogen granule protein 16 (ZG16) displayed enlarged abdominal fat pats, and these enlarged fat pads were not observed when the ZG16^-/-^ mice were depleted of their microbiota (Bergström et al., 2016). Consequently, a defective mucus barrier may allow the translocation of commensal intestinal bacteria across the epithelial barrier, which may eventually lead to metabolic impairments (Stenman et al., 2016).

In conclusion, we here identified that a previous – but not recent – history of antibiotic use in humans shapes a microbial community that is insufficient to maintain proper mucus function in mice. However, despite the usage of human-derived microbiota, further studies are required to verify that repeated antibiotic use causes similar microbiota-mediated mucus defects in humans.

## DATA AVAILABILITY STATEMENT

Human stool samples used in this study have been collected for a previously published study (Aasmets et al., 2022) and corresponding shotgun metagenomic data have been deposited previously in the European Genome-Phenome Archive database (https://www.ebi.ac.uk/ega/) under accession code <u>EGAS00001008448</u>. Due to the sensitive nature of the human phenotype data, access is restricted and must be requested through the Estonian biobank. Such access must follow informed consent regulations set out by the Estonian Committee on Bioethics and Human Research (https://genomics.ut.ee/en/content/estonian-biobank). Preliminary requests for phenotype and raw metagenome data access must be sent to.

The shotgun metagenomic sequencing data from mice and human pooled samples (human reads removed) are deposited in the European Nucleotide Archive with accession number PRJEB72415 (https://www.ebi.ac.uk/ena/browser/view/PRJEB72415).

## ETHICS STATEMENTS

### Ethics approval

Animal experiments performed at Umeå University, Sweden, were approved by the local animal ethical committee (Dnr A14-2019). The activities of the Estonian Biobank (EstBB) are regulated by the Human Genes Research Act, which was adopted in 2000 specifically for the operations of the EstBB. This study was approved by the Research Ethics Committee of the University of Tartu (approval No. 266/T10) and by the Estonian Committee on Bioethics and Human Research (Estonian Ministry of Social Affairs; approval No. 1.1-12/17).

## Acknowledgements

The authors thank the Estonian Biobank research team (Mait Metspalu, Andres Metspalu, Lili Milani, Tõnu Esko) from the Estonian Genome Centre, Institute of Genomics, University of Tartu, for collection of the health records data for the EstBB. Services from the Biochemical Imaging Centre Umeå (BICU) as part of the National Microscopy Infrastructure NMI (VR-RFI 2019-00217), Umeå Hypoxia Research Facility (UHRF), the Umeå Plant Science Centre (UPSC) Microscopy Facility, Umeå Centre for Comparative Biology (UCCB) and the FIMM Metabolomics/ Lipidomics/ Fluxomics Unit (Finland), supported by HiLIFE, are further acknowledged.

## Author contributions

Conceptualisation, KLK, OA, RHF, TT, TO, EO, BOS; Methodology, KLK, RHF, SW, SMH, FPB, SW, KP, OA, BOS, Investigation, KLK, RHF, SW, SMH, FPB, RA, KP, EO, BOS; Writing – Original Draft, KLK; Writing – Review & Editing, all authors; Funding Acquisition, TT, BOS, EO; Resources, BOS, EO; Supervision, EO, BOS.

---

**Supplementary Figure 1.** Absolute quantification of expression levels of tight junction proteins (ZO-1, Claudin-1), housekeeping gene (GAPDH), and inflammatory markers (TNF-α, IL-1β).

**Supplementary Figure 2.** Anaerobic BHI plate CFU counts from mouse stool for each time point during the Faecal Microbiota Transplant (FMT) experiment. BL – mouse baseline, WO1 – 1^st^ washout, WO2 – 2^nd^ washout, FMT-C – mice that will receive FMT from humans with no history of antibiotic use in 10 years preceding stool sample collection, FMT-ABX – mice that will receive FMT from human donors with a history of repeated antibiotic use.

**Supplementary Figure 3.** Non-significant correlations between SCFA peak area and CLR of top abundant genera in mouse gut following Faecal Microbiota Transplant (FMT). (A) Acetate correlations; (B) Butyrate correlations; (C) Propionate correlations; (D) Valerate correlations. FMT-C – mice that received FMT from humans with no history of antibiotic use in 10 years preceding stool sample collection. FMT-ABX – mice that received FMT from human donors with a history of repeated antibiotic use. P-values correspond to Spearman or Pearson correlation, as noted, depending on normality distribution of the data, p-value correction was performed using the Benjamini-Hochberg procedure. * - p < 0.05.

