## Supplementary Tables 1 to 3 for "A history of repeated antibiotic usage leads to microbiota-dependent mucus defects"

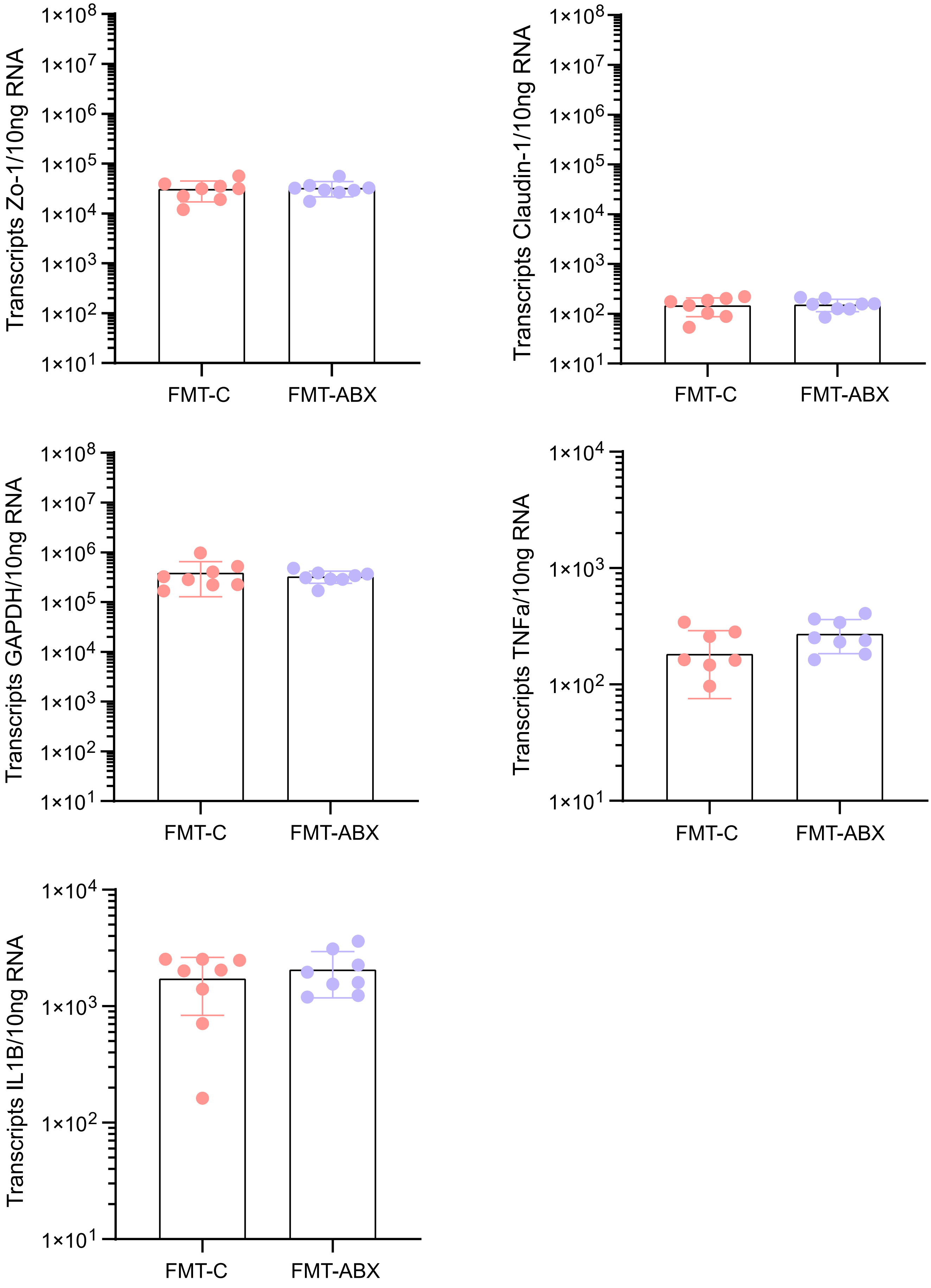


**Supplementary Figure 1.** Absolute quantification of expression levels of tight junction proteins (ZO-1, Claudin-1), housekeeping gene (GAPDH), and inflammatory markers (TNF-α, IL-1β).

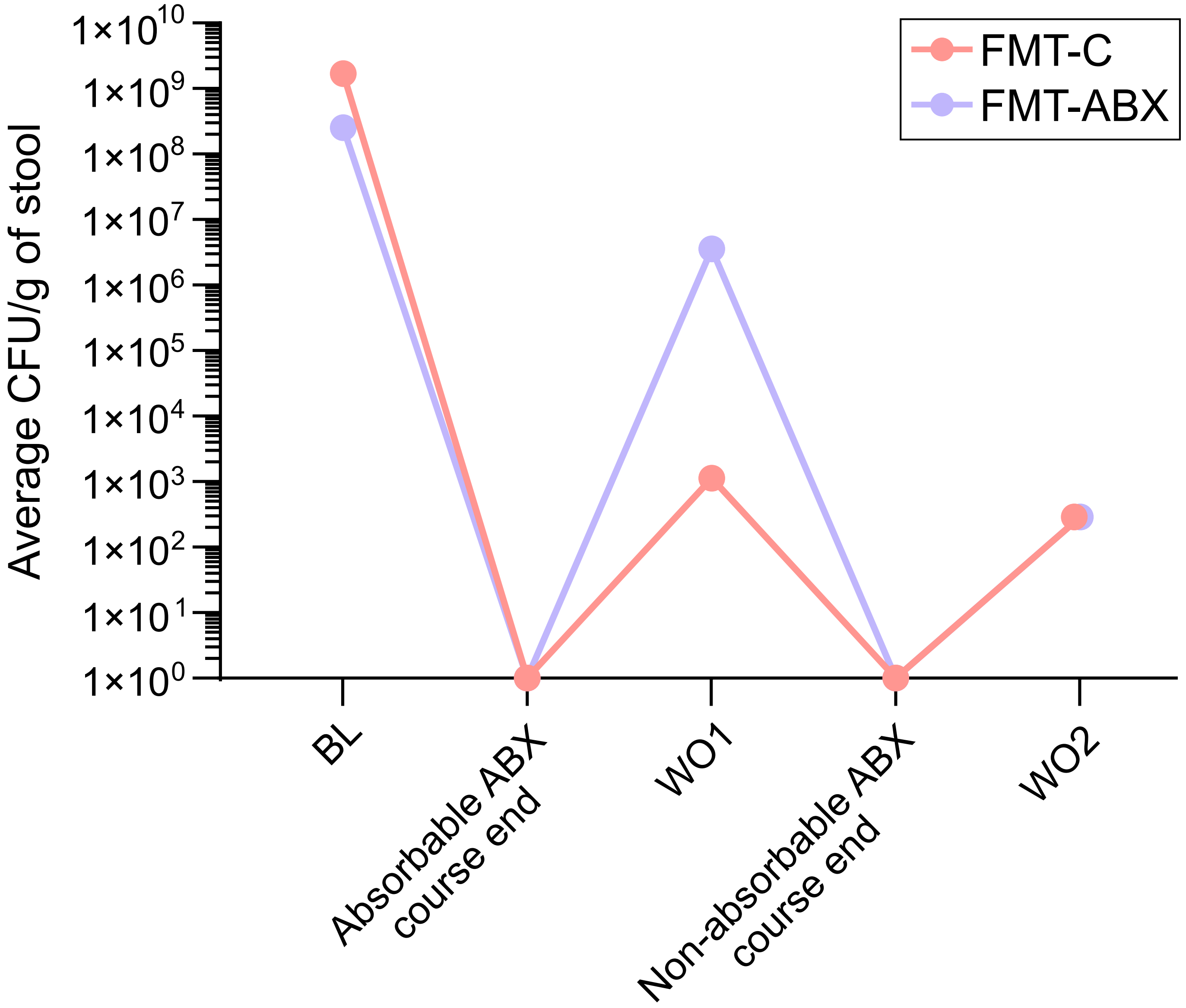


**Supplementary Figure 2.** Anaerobic BHI plate CFU counts from mouse stool for each time point during the Faecal Microbiota Transplant (FMT) experiment. BL – mouse baseline, WO1 – 1st washout, WO2 – 2nd washout, FMT-C – mice that will receive FMT from humans with no history of antibiotic use in 10 years preceding stool sample collection, FMT-ABX – mice that will receive FMT from human donors with a history of repeated antibiotic use.

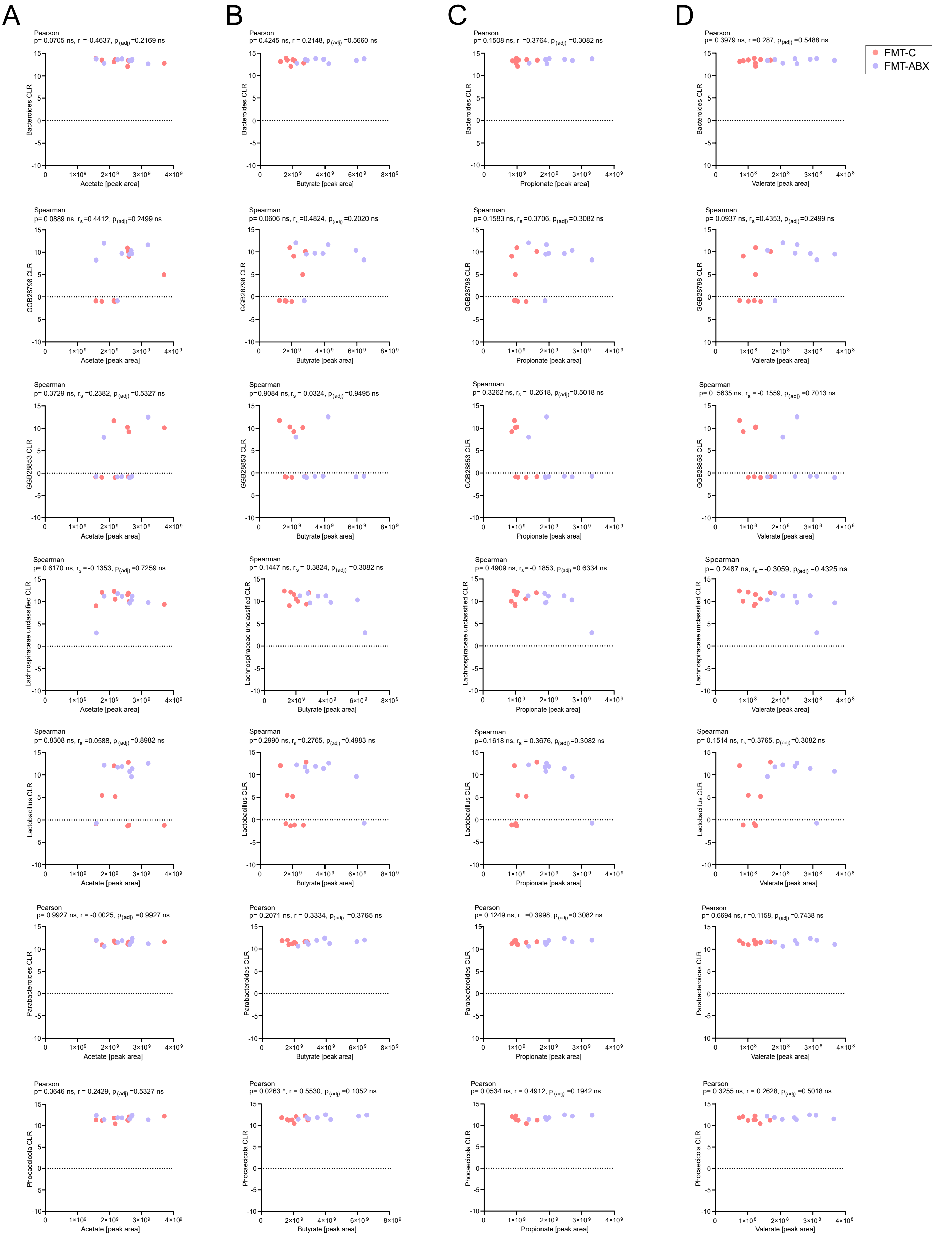


**Supplementary Figure 3.** Non-significant correlations between SCFA peak area and CLR of top abundant genera in mouse gut following Faecal Microbiota Transplant (FMT). (A) Acetate correlations; (B) Butyrate correlations; (C) Propionate correlations; (D) Valerate correlations. FMT-C – mice that received FMT from humans with no history of antibiotic use in 10 years preceding stool sample collection. FMT-ABX – mice that received FMT from human donors with a history of repeated antibiotic use. P-values correspond to Spearman or Pearson correlation, as noted, depending on normality distribution of the data. P-value correction was performed using the Benjamini-Hochberg procedure. * - $p < 0.05$.
